# PACS allows comprehensive dissection of multiple factors governing chromatin accessibility from snATAC-seq data

**DOI:** 10.1101/2023.07.30.551108

**Authors:** Zhen Miao, Jianqiao Wang, Kernyu Park, Da Kuang, Junhyong Kim

## Abstract

Single nucleus ATAC-seq (snATAC-seq) experimental designs have become increasingly complex with multiple factors that might affect chromatin accessibility, including genotype, cell type, tissue of origin, sample location, batch, etc., whose compound effects are difficult to test by existing methods. In addition, current snATAC-seq data present statistical difficulties due to their sparsity and variations in individual sequence capture. To address these problems, we present a zero-adjusted statistical model, Probability model of Accessible Chromatin of Single cells (PACS), that can allow complex hypothesis testing of factors that affect accessibility while accounting for sparse and incomplete data. For differential accessibility analysis, PACS controls the false positive rate and achieves on average a 17% to 122% higher power than existing tools. We demonstrate the effectiveness of PACS through several analysis tasks including supervised cell type annotation, compound hypothesis testing, batch effect correction, and spatiotemporal modeling. We apply PACS to several datasets from a variety of tissues and show its ability to reveal previously undiscovered insights in snATAC-seq data.

## Main

Single nucleus ATAC-seq (snATAC-seq) is a powerful assay for profiling the epigenetic status of open chromatin in individual cells^1,2^, and has been applied to study gene regulation across tissues and under various conditions, including homeostasis^3,4,5^, development^6,7^ or disease^8,9^. The cis-regulatory elements (CREs), modulated by nucleosome turnover and occupancy^10^, display variable accessibility across cells. In a cell, the dynamic of CREs activities are dependent on various physiological factors such as cell type^1,3^, developmental state^6,7^, spatial location of the tissue^11,12^, as well as interaction of these factors with genetic variation^13,14^. Identifying the sets of elements whose accessibility is governed by genetic, developmental, and physiological factors is essential in understanding the cis-regulatory codes of biological processes^15,16^. Moreover, prioritizing regulatory element is instrumental for identifying relevant cell types associated with complex traits^5,17–20^. Dissecting genetic, epigenetic, and environmental factors requires analysis methods that allows compound hypothesis testing. However, the discrete and sparse nature of ATAC-seq data presents technical challenges for existing approaches. Here, we establish a model framework and a new statistical method that allows complex hypothesis testing for single nucleus ATAC-seq data.

Among all the factors that drive the accessibility of CREs, only some factors are experimentally controlled. In a typical single cell experiment, the collection of cells is a random sample of a cell’s variable states over the unknown factors (e.g., cell cycle stage, metabolic cycles) while controlling for the known factors (e.g., tissue, location, batch). We note that sometimes the values of these factors are estimated from the data, such as unsupervised inference of cell type labels or time-sequences. Nevertheless, as the data are sampled over unknown microstates and stochastic molecular processes, the latent accessibility of a CRE should be considered as a random variable, even without experimental variability.

A central theme in snATAC-seq data analyses is to understand the dynamic of CRE across cell types and biological conditions. This is typically achieved by conducting a statistical test for Differential Accessible Regions (DARs) between different conditions of hypothetical causal factors that might govern the chromatin structure (e.g., treatment vs control). However, most existing approaches for identifying DARs rely on pairwise testing of one putative causal factor at a time (e.g., ArchR^21^, snapATAC^22^, or contingency table-based approaches) that does not allow for simultaneously testing multiple factors or testing continuous variables (such as temporal effects). A common practice in literature is to ignore other covariates while testing the factor of interest. However, as schematically diagramed in Fig. 1, testing one factor at a time can create false negatives and false positives in DAR of main factors and cannot test for interactions. Although Seurat/Signac^23,23^ uses the logistic regression framework which, theoretically, can conduct multi-factor testing, its current implementation creates spurious dependency problems that result in mis-calibrated tests (**Supplemental Note 1**).

**Figure 1.**
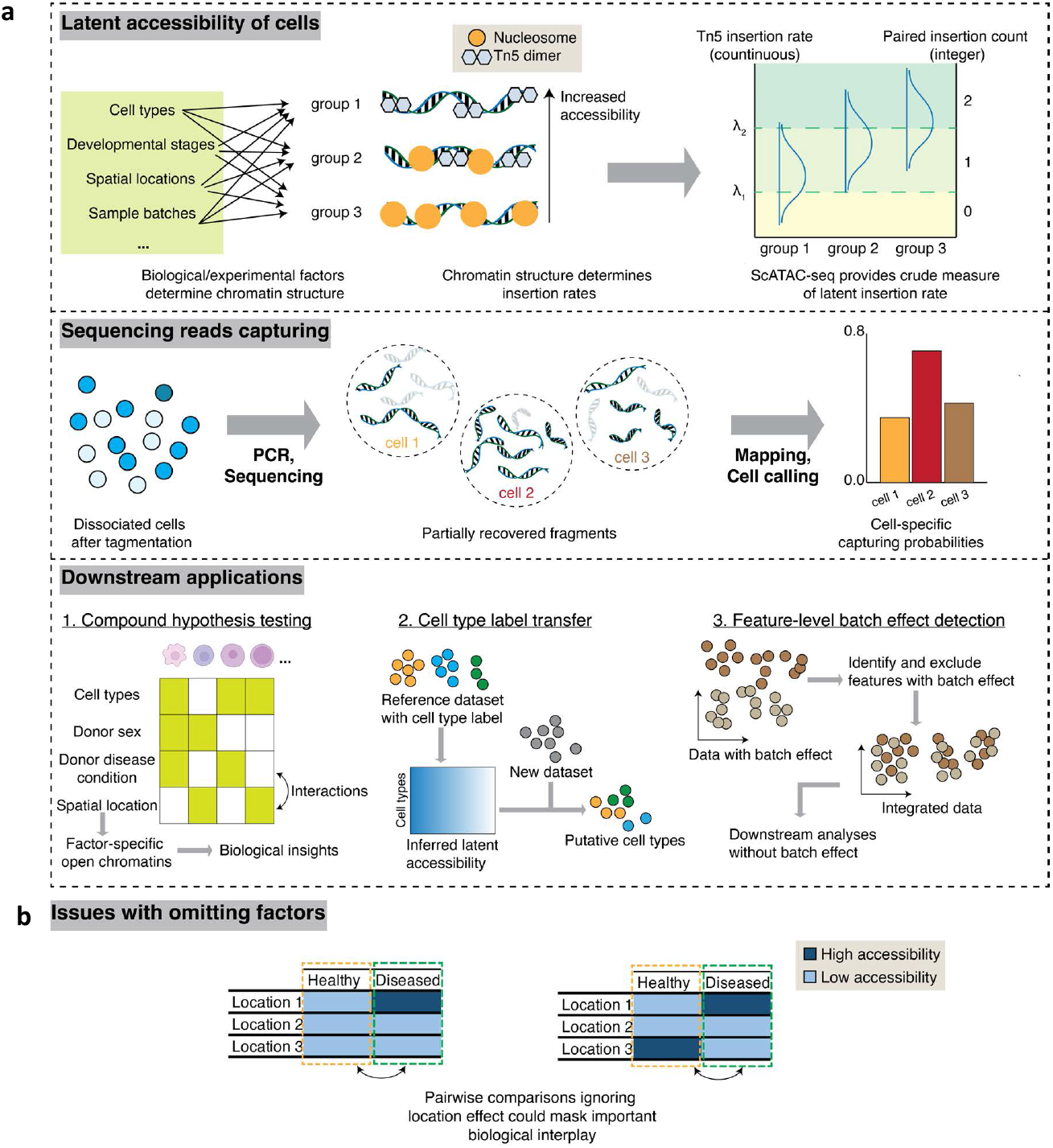
PACS modeling framework. **a**. Upper panel: Illustration of the latent accessibility of cells. Multiple factors including cell types, developmental stages, spatial locations etc. determines the chromatin structure and configurations of corresponding cell groups. These different chromatin structures result in the variable Tn5 insertion rates in the ATAC-seq experiments. The readout of ATAC assays are paired insertion counts (PIC), which are crude measures of latent insertion rates. Middle panel: Illustration of the sequencing reads capturing process of snATAC-seq. During PCR and sequencing, fragments in each single cell are partially captured, and after data processing, variable capturing probability should be accounted for in data modeling. Lower panel: Example downstream application of PACS. PACS provide functionalities for compound hypothesis testing, cell type label transfer, and feature-level batch effect correction. **b**. Example chromatin configurations where ignoring factors in conducting Differential Accessible Regions (DARs) test may result in false interpretation.

Besides the key problem of inability to hypothesis test multiple causal factors together, there are two additional unresolved technical problems: (1) snATAC-seq data displays heterogeneous sequencing coverage in each cell, so the cells are not directly comparable; (2) snATAC-seq data is sparse but also contains quantitative information at each peak^24^. To address these challenges, we present a new statistical framework, missing-corrected cumulative logistic regression (mcCLR) for the analysis of snATAC-seq data with modified Firth regularization^25,26^ to account for data sparsity.

With this statistical framework, we present our Probability model of Accessible Chromatin of Single cells (PACS), a toolkit for snATAC-seq analysis. PACS allows for complex compound analysis tasks in snATAC-seq data, including cell type classification, feature-level batch effect correction, and spatiotemporal data analysis. With simulated data and real data, we show that PACS effectively controls false positives while maintaining sensitivity for model testing. We apply PACS to a mouse kidney dataset, a developing human brain dataset, and a time-series PBMC treatment dataset, all of which have complex study designs, to demonstrate its capability to model multiple sources of variations for hypothesis-driven biological inference.

## Results

### Probabilistic model of accessible peaks and statistical test framework

In the PACS framework, we model the accessibility state of CREs in a single cell as a function of predictive factors such as cell type, physiological/developmental time, spatial region, etc. We use a matrix *F*_*C*×*J*_ to represent these variables in each cell, where *C* is the number of cells and *J* is the number of predictive variables (including dummy variables). Let *Y*_*C*×*M*_ represent an integer-valued snATAC-seq count matrix across *C* cells and *M* genomic regions. For empirical ATAC-seq data, these regions *M* are determined by data-dependent peak calling, where peaks are regarded as the set of candidate CREs^27,28^. As snATAC-seq can recover quantitative information on the density and distribution of nucleosomes^24,29^, we use integer values *Y*_*cm*_ ∈ {0,1,2, …} to represent the level of accessibility. Existing pipelines diverge in the quantification of snATAC-seq counts, and we propose to use the paired insertion count (PIC) matrix as a uniform input for downstream analyses^24^. For standard snATAC-seq experiments, PIC counts follow a size-filtered signed Poisson (ssPoisson) distribution for a given Tn5 insertion rate^24^. Thus, the integer-valued PIC counts are observed measurements of the latent Tn5 insertion rates and chromatin accessibility (**Fig. 1**, upper panel). Based on this latent variable perspective we developed a proportional odds cumulative logit model to decompose the cumulative distribution of *Y*_*cm*_ by its predictive variables *F*_*C*\*_.

With cell-specific nucleosome preparation and sequencing depth, the (observed) snATAC-seq output may miss sequence information from certain accessible chromatin (**Fig. 1**, middle panel). Here, we use *R*_*C*×*M*_, with binary values, to represent the read recovery/capturing status for each cell and region. This matrix encapsulates all the experimental factors (Tn5 activities, sequencing depth, etc.) that result in a disparity of reads recovered across cells. The observed chromatin states, denoted by *Z*_*CM*_, are specified by the element-wise product between the latent accessibility *Y*_*CM*_ and the capturing status *R*_*CM*_. Since various experimental factors such as sequencing depth are cell-specific, we further assume the capturing probability *P*(*R*_*CM*_ = 1) to be unique to each cell but common to all peaks in that cell, and thus we use *q*_*c*_ to denote this conditional read capturing probability in cell *c*.

Motivated by the latent variable model and to account for cell-specific missing data, we extended the cumulative logit model to simultaneously decompose accessibility as:

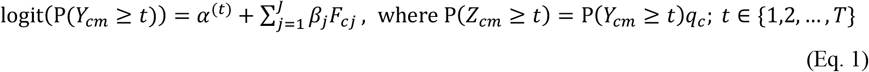

where *q*_*c*_ is the capturing probability for a cell *c, P*(*Y*_*cm*_ ≥*t*) is the sampling probability of cells with accessibility level greater than or equal to *t, α*^(*t*)^ is the intercept term in the *t*^*th*^ cumulative logit, and *β*_*j*_ is the coefficient for the *j*^*th*^ column of the design matrix. Eq. 1 assumes a proportional odds model, where we have a common set of coefficients *β*_*j*_ for all levels of the cumulative distribution, while allowing for a unique constant term *α* ^(*t*)^ for each level. Hereafter, we refer to our method as the **mcCLR** model, which stands for the missing-corrected cumulative logit regression model.

With the formulation above, the effect of a complex set of predictive variables (and their interactions) on accessibility can be tested by the null hypothesis of *β*_*i*_ = 0 with a likelihood ratio test (**Fig. 1**, lower panel). One statistical challenge is to estimate *q*_*c*_’s for each cell. We assumed the same capturing probability within a cell regardless of accessibility across different peaks such that the problem is tractable and can be computed efficiently. Operationally, we first group the cells by their combination of the treatments and then utilize a coordinate descent algorithm to obtain estimates of *P*(*Y*_*cm*_ ≥ 1|*F*_*C*_) and *q*_*c*_ (**Methods**).

Another statistical challenge of snATAC-seq is that the data is very sparse, creating a so-called “perfect separation” problem (see^30^). Here, we developed a regularized model to resolve the issues with sparsity in snATAC-seq data by generalizing the Firth logistic regression model^25,31^, where we incorporate the cell-specific capturing probability (Eq. 1) into the model (**Methods**). Essentially, a Firth penalty is introduced in the regression model:

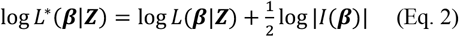

Where *L*^*^ represents the penalized likelihood, *L* is the likelihood of the regression model, and *I*(***β***) is the information matrix. Derivations of the parameter estimation framework are described in the **Methods** section. With the proposed methods, we aim to control type I error more accurately and account for technical zeros (due to uneven data capturing) and sparse data. This regression-based model enables the testing of multiple covariates that jointly determine accessibility, while controlling for other covariates or confounders.

### Application of PACS to cell type identification

To demonstrate the effectiveness of our model for separating the latent chromatin accessibility from the capturing probability, we evaluated three model assumptions using the task of (supervised) cell type prediction, where the goal is to predict cell types in a new snATAC-seq dataset given an annotated (labeled) dataset.

We first evaluated the accuracy of the estimation procedure of PACS. We simulated groups of cells with a spectrum of both the underlying probability of accessibility (*P*(*Y*_*cm*_ ≥ 1), or *p* in short) across peaks, and the capturing probabilities (*q*) across cells (**Methods**). We then utilized PACS to jointly estimate *p* and *q*, with n=1000, 500, or 250 cells. The simulation results show that our estimator can determine both the capturing probabilities and open-chromatin probabilities accurately, with root mean squared errors (RMSE) for the underlying probability of accessibility from 0.028 (n=1000) to 0.027 (n=250) and RMSE for capturing probability from 0.0067 (n=1000) to 0.012 (n=250, **Supplementary Fig. 1a-f**, and **Supplementary Table 1**).

**Table 1.**
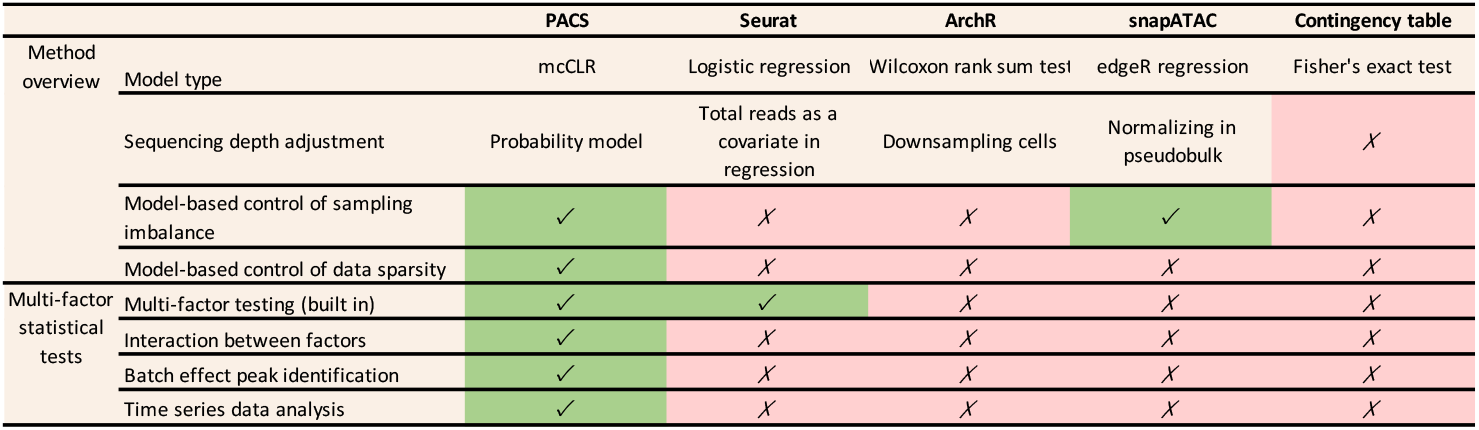
Comparison of model design and capability among existing scATAC-seq testing tools.

We next tested PACS by applying it to a cell type label transfer task, comparing it with the Naïve Bayes model. For both models, we started with an estimated *P*_*g*_ for each known cell type group label *g*, and then applied the Bayes discriminative model to infer the most probable cell type labels for novel unidentified cells. Naïve Bayes does not assume missing data; thus, it ignores the cell-specific capturing probability. The prediction performances were evaluated with ten-fold cross-validation and holdout methods, where the original cell type labels are regarded as ground truth (**Methods**). We tested the methods on five datasets, including two human cell line datasets^21^, two mouse kidney datasets^6^, and one marmoset brain dataset^32^. In the two human cell line datasets, the cell line labels are annotated by their SNP information^21^, so the labels are regarded as gold standards. For the remaining datasets, the original cell type labels are generated by clustering and marker-based annotation, so the labels may have errors.

PACS consistently outperforms the Naïve Bayes model with an average 0.31 increase in Adjusted Rand Index (ARI, **Supplementary Fig. 2a**), suggesting the importance of considering the cell-to-cell variability in capturing rate. For the gold-standard cell line mixture data, we achieved almost perfect label prediction (ARI > 0.99), while Naïve Bayes had much lower accuracy with an average ARI = 0.54 (**Supplementary Fig. 2b-c**). For the kidney data^6^ and the marmoset brain data^32^, PACS still achieved high performance, with average ARI equal to 0.92, 0.90, and 0.88 for the adult kidney, P0 kidney, and marmoset brain data, respectively. The Naïve Bayes model, on the other hand, again produced lower ARI scores, equal to 0.59, 0.65, and 0.69 for the three datasets, respectively (**Supplementary Fig. 1d-k**).

**Figure 2.**
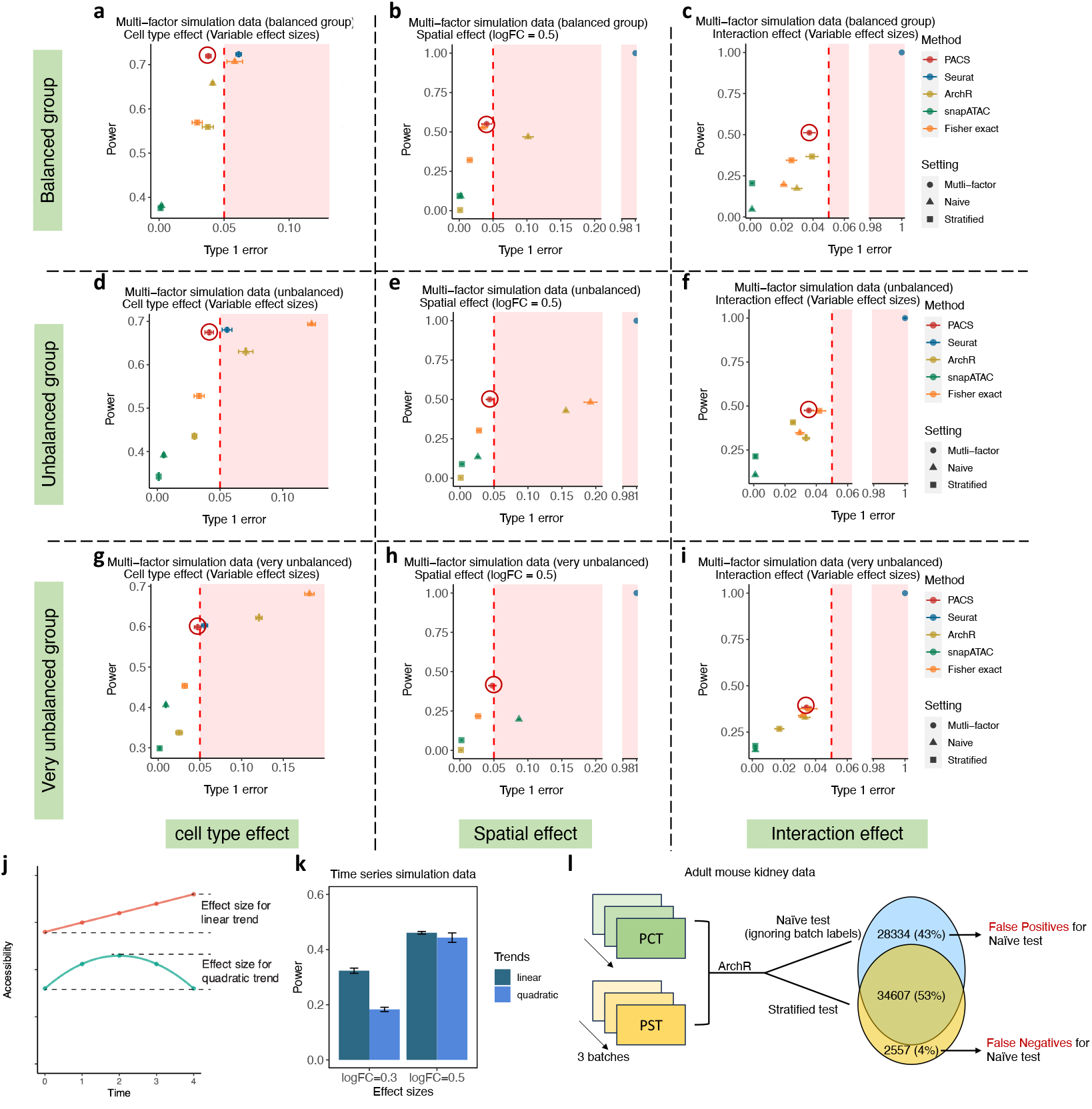
Compound hypothesis testing with PACS is sensitive and specific. **a-i**. Type I error and power of different methods evaluated with two-factor simulation data. All tests are two-sided. The best-performing method is indicated by a circle with matched color. **j**. Illustration of linear and quadratic effects of treatment on accessibility across five time points. Effect sizes are defined as the log fold change (logFC) between the highest accessibility over the lowest accessibility. **k**. Evaluation of power in detecting linear and quadratic temporal effects using simulated data with different effect sizes. N=1000 cells were sampled for each time point. **I**. The false positive and false negative rates from ignoring batch covariates in testing cell type effect with ArchR, for the adult mouse kidney data.

For the holdout experiment, where training and testing is done on different datasets, consistent with the above results, our method shows more accurate cell label prediction than Naïve Bayes. We note that our cell type label prediction approach is very efficient, and the total time for training and prediction takes < 5 min for large datasets (>70,000 cells).

### PACS enables parametric multi-factor model testing for accessibility

Identifying the set of CREs regulated by certain physiological cues is essential in understanding functional regulation. For example, differentially accessible region (DAR) analysis tries to determine cell type-specific chromosomal accessibility differences. Most snATAC-seq pipelines adopt RNA-seq differential expression methods to ask whether a peak belongs to a DAR. These approaches generally lack calibration for sparse ATAC data, and the approach of pairwise DAR tests does not allow testing more complex models that might determine peak accessibility (e.g., combination of spatial location, batch effects). With existing methods for DAR detection, commonly adopted approaches are to ignore other factors or stratify by other factors to test the factor of interest, if the predictive variables are nominal (e.g., cell types). However, such tests involve ad hoc partition into levels of the nominal factor and cannot test more complex models including possible metric variables (e.g., developmental time).

Here we first assess model design and capabilities of PACS and four established tools/methods: ArchR^21^, Seurat/Signac^23^, snapATAC^22^, and Fisher’s exact test. ArchR conducts Wilcoxon rank-sum test on the subsampled cells from the initial groups, while ensuring parity in the number of sequencing reads across any two samples being tested. Seurat employs a standard logistic regression model^33^, but setting the tested factor as the dependent variable and read counts (Tn5 insertions) and total reads as predictive variables. SnapATAC conducts a test on the pseudo-bulk data of two groups and utilizes the edgeR^34^ negative binomial test on the pseudo-bulk data with a pre-defined ad hoc variance measure (biological coefficient of variation, bvc = 0.4 for human and 0.1 for mouse data). A comprehensive comparison is summarized in **Table 1**.

To quantitatively compare the performance of the parametric test framework, we first used simulated data to test a single factor model (cell types as factor). To resemble real data, simulated samples were generated by parameterizing the model with the accessibility and capturing probability estimated directly from a human cell line dataset^21^ (**Methods**). We randomly sampled varying numbers of cells in each group, ranging from 250 to 1000. **Supplementary Fig. 3** shows that Seurat failed to control type I error at the specified significance level. Among the methods demonstrating type I error control, PACS has on average 17%, 19% and 122% greater power than Fisher’s exact test, ArchR and snapATAC, respectively (**Supplementary Table 2**). The reduced power of ArchR is likely due to the subsampling process, and the ad hoc “bvc” choice in snapATAC may result in a miscalibrated test with a low type I error and power. The q-q plots of the five methods are shown in **Supplementary Fig. 4a-e**.

**Figure 3.**
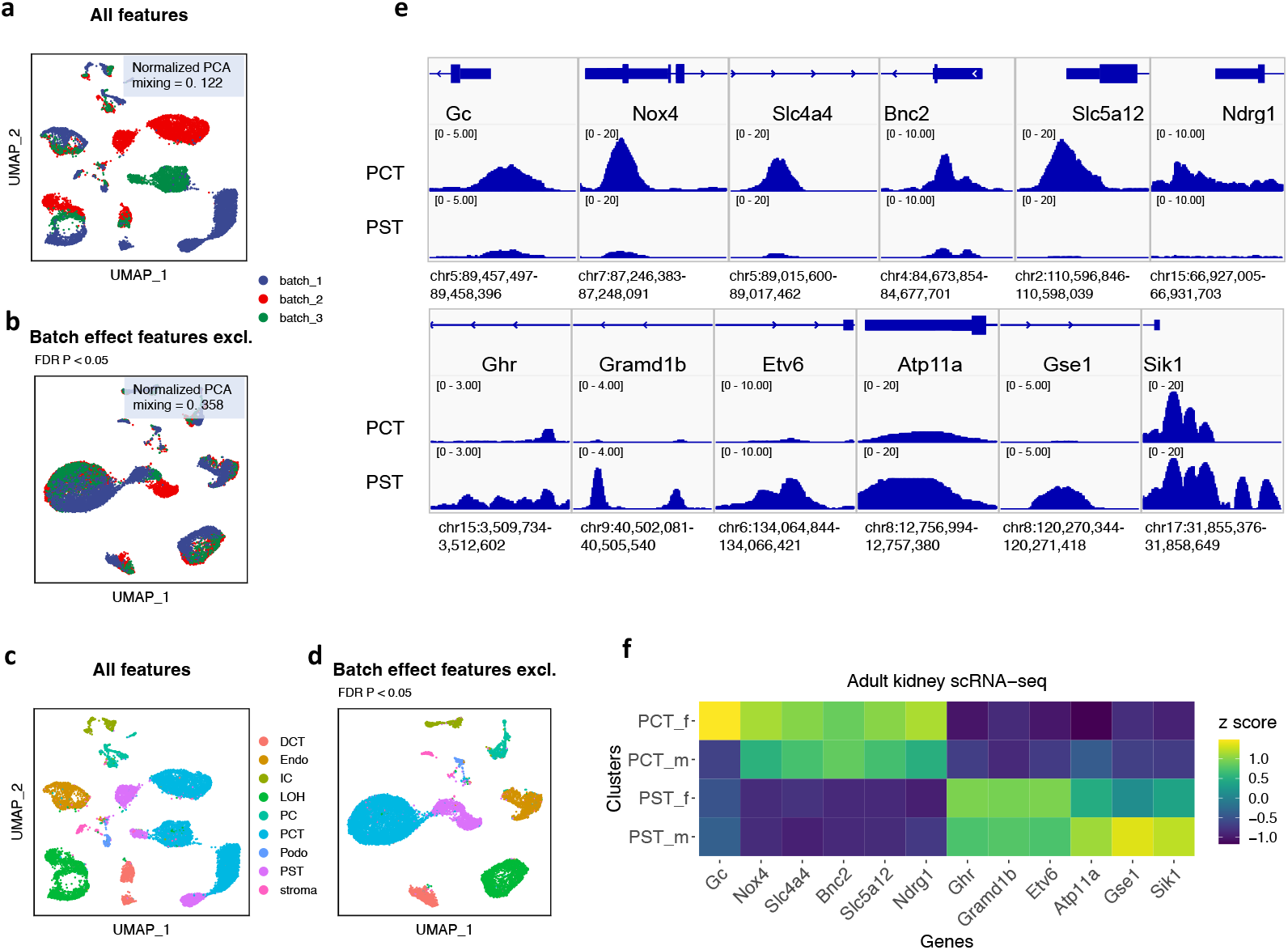
Application of PACS to the mouse kidney dataset. **a-b**. UMAP dimension reduction plot constructed with all features (**a**) or after excluding features with significant batch effect (**b**), colored by batch labels. Features with batch effect are detected with PACS differential test module, and FDR multiple testing correction is applied. Normalized PCA mixing represents the normalized mixing score calculated in the PCA space, with 1 being no batch effect and 0 being strongest batch effect. **c-d**. UMAP dimension reduction plot constructed with all features (**c**) or after removing features with batch effect (**d**), colored by cell types. **e**. IGV plot of peak summits around cell type-specific genes identified by PACS, for PCT and PST cell types. The list of cell type specific genes is generated with GREAT enrichment analysis using differentially accessible peaks. **f**. Heatmap of normalized gene expression z scores for the scRNA-seq data from male (-m) and female (-f) kidneys. The list of genes match those from the panel **f**.

**Figure 4.**
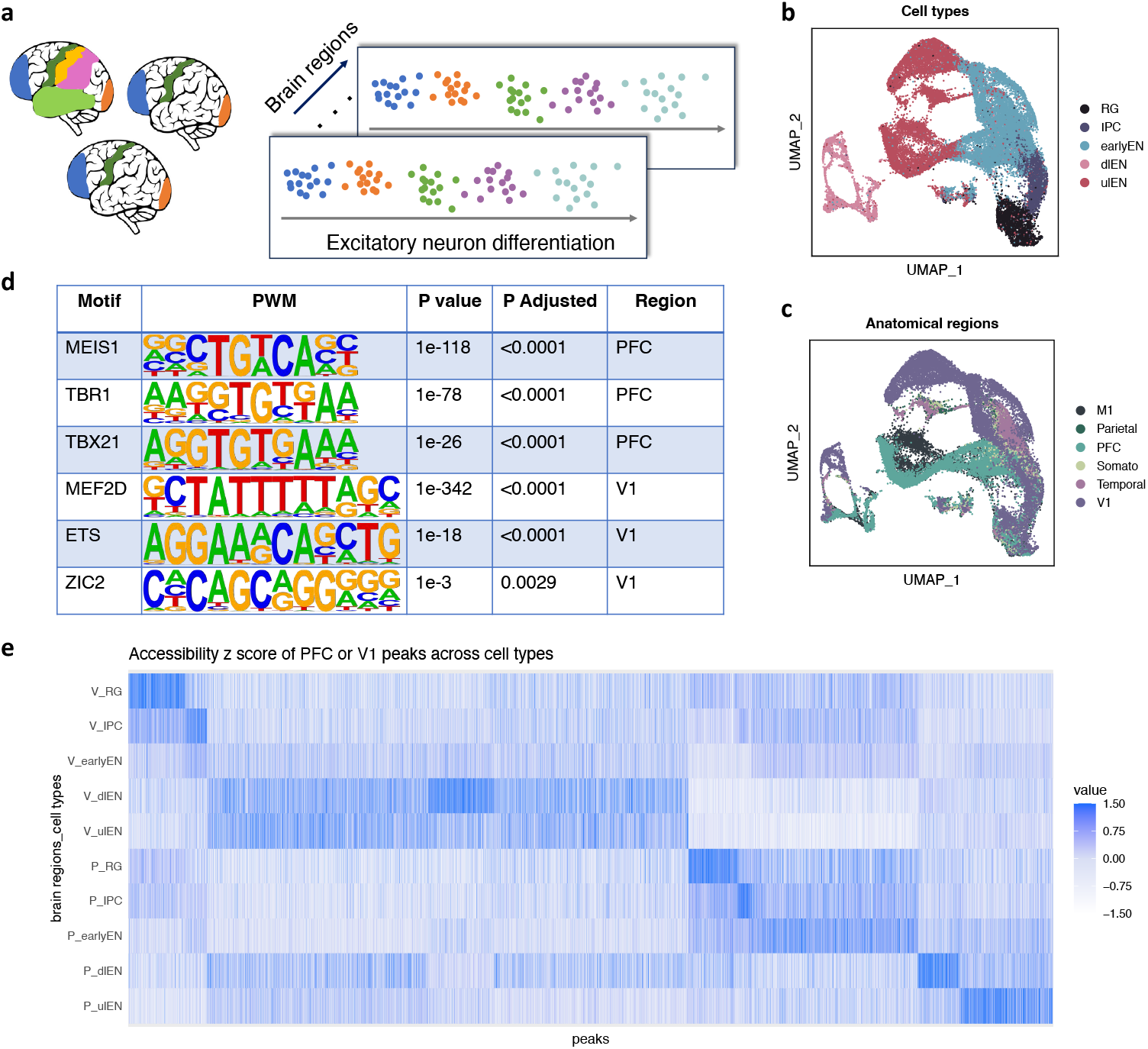
Application of PACS to the developing human brain data. **a**. Illustration of the developing human brain dataset. The subset of data we analyzed are composed of samples from three donors across six brain anatomical regions, and we focused on the excitatory neuron lineage. **b-c**. UMAP visualization of the data complexity, with points colored by cell type (**b**) or anatomical regions (**c**). RG, radial glia; IPC, intermediate (neuro-) progenitor cells; earlyEN, early excitatory neurons; dlEN, deep layer excitatory neurons; ulEN, upper layer excitatory neurons; M1, primary motor cortex; Parietal, dorsolateral parietal cortex; PFC, dorsolateral prefrontal cortex; Somato, primary somatosensory cortex; Temporal, temporal cortex; V1, primary visual cortex. **d**. Motif enrichment results for PFC- and V1-specific peaks identified using PACS. PWM, position weight matrix. **e**. Accessibility z score of PFC and V1 peaks across five cell types.

To evaluate the performance under a multi-factor model, we next conducted another set of simulations with a second factor, two spatial locations (S1 and S2) along with the first factor of two cell types (T1 and T2). We evaluated the performance in testing for the main effects or their interactions (**Methods**). For the methods that cannot directly test effects for multiple factors, two strategies were used. The first is called the “naïve test”, where the other factor is ignored in testing the factor of interest. The second strategy is called the “stratified test”, where we stratified the dataset by the second factor and conducted a pairwise test between the factor of interest on each stratum, followed by *P* value combination test (**Methods**). Across all methods and test strategies, only PACS, snapATAC (naïve and stratified), Fisher-stratified, and ArchR-stratified controlled type I error at the specified level (**Fig. 2a-i**); PACS remained the most powerful test and detected up to 66% more true differential peaks compared with the second most powerful methods (**Supplementary Table 3**). Notably, Seurat could not determine peaks with spatial effects or interaction effects, because the cell type factor is treated as the dependent variable in the regression model. More detailed explanation can be found at the **Supplementary Note 1**.

We then simulated a time-series dataset with five time points, to evaluate our model performance for ordinal covariates. We assumed two temporal trends of accessibility, linear and quadratic trends. To put this in a biological setting, the quadratic trend may represent the presence of an acute spike response and the linear trend may represent temporally accumulating chronic responses. The PACS framework could detect both linear and quadratic signals, and its power is dependent on the “effect sizes” defined as the log fold change of accessibility between the highest and lowest accessibility (**Fig. 2j-k**).

We also evaluated the PACS model in real datasets. As the ground truth is unknown, we utilized a sampling-based approach. We used randomly permuted cell type labels to estimate the type I error. To evaluate power, we treated the consensus DAR set from all methods as “true DARs” (after type I error calibration, see **Methods**). For the standard two-group DAR test, our method consistently controlled type I error and achieved high power, across different datasets (**Supplementary Fig. 5a-f**). We further showed that ignoring confounders such as batch effect could result in substantial false discoveries, up to 47% in the adult mouse kidney data (**Fig. 2l**).

**Figure 5.**
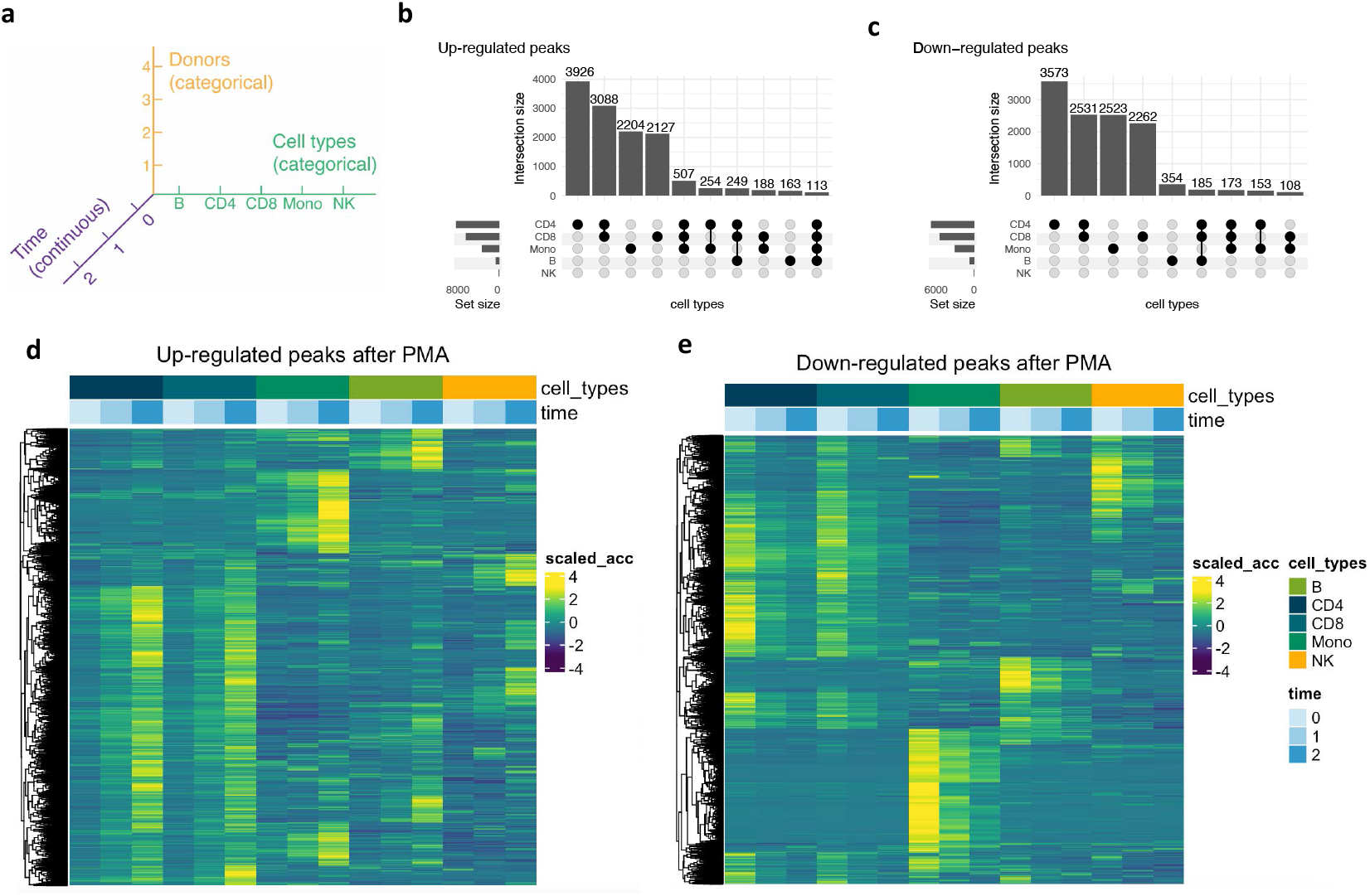
Application of PACS to time-series dataset from human PBMC treatment data. **a**. Factor landscape of the PBMC treatment dataset. Here, another layer of factor is the four different treatments, which can also be jointly considered in the model, but for demonstration purposes, we only focus on the PMA treatment effect. The control time point is considered as time 0, and the times of one and six hours after treatment are considered as time 1 and 2, respectively. **b-c**. Summary of significant up- or down-regulated peaks after PMA treatment for each cell type. **d-e**. Heatmap of significant up- or down-regulated peaks after PMA treatment, grouped by time point and cell type. The color scale (scaled_acc) represents the accessibility z score.

### PACS identifies kidney cell type-specific regulatory motifs and allows direct batch correction

One important feature of PACS is its ability to handle complex datasets with multiple confounding factors. To test the performance of PACS, we analyzed an adult kidney dataset with strong batch effects^6^. This dataset contains three samples generated independently (in three batches), and the authors identified a strong batch effect. Existing methods for batch correction map the ATAC-seq features to a latent vector space to subtract the batch effects. For example, the original study^6^ relies on Harmony^35^ to remove the batch effect in latent space for visualization and clustering, but the batch effect is still present in the peak feature sets, which could confound downstream analyses and inferences.

To remove the batch effect at the feature level, we assume that the batch effect will affect (increase or decrease) the accessibility of certain peaks, and these effects are orthogonal to the biological effects. This assumption is necessary for most of the existing batch-effect correction methods (e.g., MNN^36^, Seurat^37^, and Harmony^35^), as a matter of experimental design. With this assumption, we applied PACS on the adult kidney data, detected significant DAR peaks among batches (*P* value < 0.05 with or without FDR correction) and removed batch-effect peaks from the feature set. We next implemented Signac to process the original data as well as the batch effect-corrected data, without any other batch correction steps. Dimension reductions with UMAP suggested that the original data contained a strong batch effect, where almost all cell types are separated by batch (**Fig. 3a-b**). After removing the peaks with strong batch effects, the cells are better mixed among batches (**Fig. 4c-d, Supplementary Fig. 6a-b**). Note that different cell types are still separated, suggesting the biological differences are (at least partially) maintained. Since UMAP visualization may not fully preserve the actual batch mixing structure, we adopted a batch mixing score from Ref.^38^ to quantify the batch effect in the PCA space. The batch mixing score is defined as the average proportion of nearest neighbor cells with different batch identities, where a higher score indicates better mixing between batches, and thus a smaller batch effect (**Methods**). We normalized the mean batch mixing score by dividing it by the expected score under the random mixing scenario. After batch effect correction with PACS, the normalized mean batch mixing score is 0.358 (with FDR correction) or 0.417 (without FDR correction) compared with 0.122 before batch correction. Example peaks with strong batch effect detected by PACS are shown in **Supplementary Fig. 7a-l**.

We next applied our method to identify cell type-specific features while adjusting for batch effect. We focused on the two proximal tubule subtypes, proximal convoluted tubules (PCT) and proximal straight tubules (PST). By fitting our mcCLR model with cell type and batch effect, we identified 19,888 and 62,368 significant peaks for PCT and PST, respectively (FDR-corrected *P* value < 0.05, **Supplementary Tables 4-5**). The original study utilized snapATAC, which reported 23,712 and 36,078 significant peaks for PCT and PST, respectively. With the batch-corrected differential peaks, we then conducted GREAT enrichment analysis^39,40^ to identify candidate PCT- and PST-specific genes (**Supplementary Tables 6-7**). We identified *Gc, Nox4, Slc4a4, Bnc2, Slc5a12*, and *Ndrg1* genes as top PCT-enriched genes, and *Ghr, Gramd1b, Etv6, Atp11a, Gse1*, and *Sik1* as top PST-enriched genes. The associated genomic pile-up figures for the CREs of these genes are shown in **Fig. 3e**, and these findings were supported by a public scRNA-seq dataset^41^ (**Fig. 3f**).

### PACS dissects complex accessibility-regulating factors in the developing human brain

We applied our method to the human brain dataset^11^, which is more challenging due to the complex study design with cells collected from six donors across eight spatial locations. Substantial sequencing depth variations among samples has also been noticed, which further complicated the analysis (**Supplementary Fig. 8a-c**). To study how spatial locations affect chromatin structure, the original reference focused on the prefrontal cortex (PFC) and primary visual cortex (V1) regions, as they were the extremes of the rostral-caudal axis^11^. With the multi-factor analysis capacity of PACS, we conducted analyses to (1) identify the region effect, while adjusting for the donor effect, (2) identify the cell-type specific region effect.

We first examined the marginal effect of brain regions on chromatin accessibility, holding other factors constant (**Methods**). For this, we focused on a subset of three donors where spatial information is retained during data collection (**Fig. 4a-c, Supplementary Table 8**). In total, we identified 146,676 brain region-specific peaks (FDR corrected *P* value < 0.05). Between PFC and V1 regions, we identified 30,455 DAR peaks, ∼20% more compared with the original study (**Supplementary Tables 9-10)**. With the region-specific DARs, we conducted motif enrichment analysis to identify region-specific TFs. For the PFC and V1 regions, we found several signals that were consistent with the original article^11^, including PFC-specific motifs *MEIS1, TBX21*, and *TBR1*, and V1-specific motifs *MEF2B, MEF2C, MEF2A*, and *MEF2D*. Moreover, we identified additional V1-specific motifs *ETS* and *ZIC2* (**Fig. 4d**), supported by the scRNA-seq data collected from the same regions^42^. We also noticed that some neuron development-associated TFs, including *OLIG2* and *NEUROG2*, are enriched in both brain regions but with different binding sites, likely due to different co-factors that open different DNA regions. Motif enrichment results for both brain regions are reported in **Supplementary Tables 11-12**.

Next, we used PACS to examine the location effect across different cell types along excitatory neurogenesis. This corresponds to testing the interaction terms between spatial location and cell types, while adjusting for donor effect (**Fig. 4e**). The previous study reported that the chromatin status of the intermediate progenitor cells (IPC) population started to diverge between PFC and V1 regions. Consistent with the article, we identified 2773 significant differential peaks between PFC and V1 at IPC stage, 52% more than snapATAC (**Supplementary Table 13**).

In sum, we show the implementation of PACS for data with three levels of factors: donor, spatial region, and cell type. PACS can be applied to study one factor or the interaction between factors while adjusting for other confounding factors, and test results have higher power.

### PACS identifies time-dependent immune responses after stimulation

The existing methods for DAR detection rely on pairwise comparisons, and thus are not applicable to ordinal or continuous factors. One such example is the snATAC-seq data collected at multiple time points. Here, we apply PACS to a peripheral blood mononuclear cell (PBMC) dataset collected at three time points (0h control, 1h, and 6h) after drug treatment^43^. Multiple treatments have been applied separately to cells collected from four human donors. While PACS can simultaneously model all drugs and conditions, we focus on the ionomycin plus phorbol myristate acetate (PMA) treatment to demonstrate the PACS workflow. The factors included in the PACS model are shown in **Fig. 5a**, where cell type and donor effects are categorical, and the time effect is coded as an ordinal variable. Note that time can be alternatively coded as a continuous variable.

We tested the treatment effect by identifying open chromatin regions that show a gradual increase or decrease in accessibility after treatment. In total, we detected 35,356 peaks with a strong treatment effect across five broad cell types (B cell, CD4 T cell, CD8 T cell, Monocyte, and NK cell, **Supplementary Tables 14-16**). Across the cell types, CD4 and CD8 T cells show the most significant changes in chromatin landscape after treatment (**Fig. 5b-c**). This is expected, as PMA can induce T cell activation and proliferation^44^. Among the peaks with significant PMA treatment effect, most become more accessible after treatment, consistent with the activation function of the treatment. We then conducted gene enrichment analysis with GREAT^40^, where we identified several GO pathways associated with T cell activation, such as “regulation of T cell differentiation” and “regulation of interleukin-2 production” (**Supplementary Table 17**). We also identified enriched genes including *DUSP5, IL1RL1, TBX21*, and *CXCR3* (**Supplementary Table 18**), expression of which have been previously reported to be up-regulated in PMA treatment^45,46,47,48^. Notably, *DUSP5* is known to play an essential role in the immune response through regulation of NF-κB as well as ERK1/2 signal transduction^49^, and *TBX21* is an immune cell TF that also directs T-cell homing to pro-inflammatory sites via regulation of *CXCR3* expression^50^. **Fig. 5d-e** showed the cell type-specific open chromatin landscape dynamic after the PMA treatment. We noticed that some CREs respond to the treatment effect across all cell types and some CREs become activated in only certain cell types.

## Discussion

Single-cell sequencing data, characterized by uneven data capturing and data sparsity, present significant challenges in data analysis. For scRNA-seq data, data normalization has been an essential step for adjusting for uneven data capturing; however, this approach is not directly applicable to scATAC-seq data, creating a unique challenge. Our method, PACS, addresses this by jointly modeling the group-level underlying accessibility and cell-level sequencing reads capturing. When applied to cell type annotation tasks, PACS showed improved performance compared with the Naïve Bayes model, which does not consider cell-specific capturing probabilities.

The increasing volume of data across various tissue conditions necessitates atlas-level data integration to comprehend tissue dynamics. Our cell type annotation framework facilitates the transfer of cell type annotation from reference dataset to another dataset, thereby overcoming a major hurdle in integrative data analysis. A further challenge of data integration is to jointly model various factors (e.g., genotype, cell type, spatial locations) that govern cellular CRE activities. The standard GLM framework could not address the uneven data capturing in snATAC-seq data, so we developed a statistical model that extends the GLM framework to account for cell-specific missing data. Our model, the missing-corrected cumulative logistic regression (mcCLR) with regularization, enables PACS to perform multi-covariate hypothesis tests including spatial and temporal data analysis. Here we analyzed three empirical datasets from brain, kidney, and blood samples to show the utility and flexibility of our framework in large, complex datasets. Additionally, PACS shows promise in genotype-based chromatin accessibility studies, such as allele imbalance analysis or chromatin accessible quantitative trait locus (caQTL) study. Compared with existing models, PACS offers a more controlled approach for handling multiple covariates in the caQTL analysis.

We have previously derived a parametric model of the snATAC-seq read count, known as the size-filtered signed Poisson distribution (ssPoisson)^24^. In our current work, we treat the insertion rate as a latent variable and directly model the paired insertion counts (PIC) of the data with a regression-based model. This approach greatly enhanced computational efficiency. Future research will delve into the potential applications of parametric distributions. In summary, PACS allows versatile hypothesis testing for the analysis of snATAC-seq data, and its capability of jointly accounting for multiple factors that govern the chromosomal landscape will help investigators dissect multi-factorial chromatin regulation.

## Methods

### Data availability

We downloaded the following snATAC-seq datasets from public repositories: mouse kidney data^6^ (GEO GSE157079, https://www.ncbi.nlm.nih.gov/geo/query/acc.cgi?acc=GSE157079), human cell line data^21^ (GEO GSE162690, https://www.ncbi.nlm.nih.gov/geo/query/acc.cgi?acc=GSE162690), developing human brain data^11^ (GEO GSE163018, https://www.ncbi.nlm.nih.gov/geo/query/acc.cgi?acc=GSE163018), marmoset brain data^32^ (the Brain Cell Data Center RRID SCR_017266; https://biccn.org/data), human PBMC time-series stimulation data^43^ (GEO GSE178431, https://www.ncbi.nlm.nih.gov/geo/query/acc.cgi?acc=GSE178431).

### Probabilistic model of underlying open chromatin status

Here we model the activity of regulatory elements in each cell type group by the cumulative distribution of the accessibility. The underlying accessibility for a CRE is a function of nucleosome density and turnover rate. As we discuss in the main text, for a particular cell group, the chromatin state should be regarded as a random variable as they are sampled from mixtures of hidden microstates. Here, we expanded the model of accessible chromatin from Ref^24^. Briefly, let *F*_*C*×*J*_ be a design matrix that summarizes known independent variables (e.g., cell type, developmental time, sample locations, etc.) across *C* cells, *Y*_*C*×*M*_ be the underlying (latent) chromatin status across *C* cells and *M* regions, where each element represent the accessibility of a genomic region. The goal of PACS is to decompose the (complementary) cumulative distribution of *Y*_*cm*_, i.e., the series of distributions:

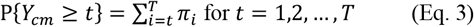

by predictive independent variables in *F*_*c*\*_. Here the maximum value of accessibility we account for, *T*, is feature specific. To be precise, for a feature *m, T* is the largest integer such that ∑_*C*_ 1(*Z*_*CM*_ ≥*t*) ≥ *n*_*C*_ where *n*_*C*_ is a hyperparameter. In our study, *n*_*C*_ is set to be 0.25*C* but based on our evaluation, our model is not sensitive to the choice of *n*_*C*_.

### Model for capturing probability of cell

Due to various experimental factors like enzyme activity and sequencing depth disparities across cells, we introduce *R*_*C*×*M*_ as a matrix representing the capturing status of each cell and region. Let *Z*_*C*×*M*_ be the (observed) scATAC dataset, we have *Z* = *Y* ⨂ *R*, where ⨂ denote element-wise Product. We consider *R*_*cm*_ to be sampled from a Bernoulli distribution parameterized by *q*_*c*_, cell-specific capturing probability:

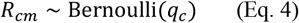

### Joint parameter estimation for single-factor scenario

Given a class of data that correspond to a combination of levels of independent variables, we follow the same parameter estimation framework as described in Ref^24^. Briefly, assume we have a genomic region-by-cell (i.e., peak-by-cell) matrix 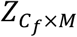 with *C*_*f*_ denoting the subset of cells corresponding to some combination of the independent prediction factors. The observed values in 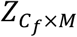 are ordinal values, but as most of the non-zero scATAC-seq counts are one (typically >70%), we focus on 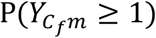 for purposes of *q*_*c*_ estimation. Hereafter, we use the notation 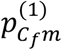 to represent the (non-zero) open probability of group *C*_*f*_ and feature *m*. We have further assumed *q*_*c*_ to be identical across different levels of accessibility for a given cell. Due to the data sparsity and the predominant counts of one, this assumption is moderate, and the estimation process will be greatly accelerated with this assumption. We use moment estimator with a coordinate descent algorithm to iteratively update 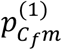 given *q*_*c*_, and update *q*_*c*_ given 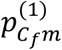. Briefly, we execute the following iteration until convergence:

1. Start with an initial estimate of 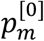
2. For *t* = 1, 2, …
  a. Compute 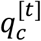 by:

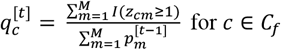
  b. Update 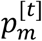 by moment estimator:

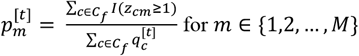

where we use superscript [*t*] to represent the *t*^*th*^ iteration, and we omit the subscript *C*_*f*_ and superscript (1) for 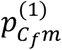.

### Uniqueness of parameter estimation

In order for the above joint parameter estimation framework to converge and for the estimated parameters to be uniquely defined, there should be *q*_*c*_ = 1 for some cells and 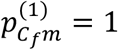 for some features. In PACS, we conduct a convergence check by requiring a certain proportion of cells (default 10%) to have an estimated capturing probability greater than 0.9. In the case of a cluster of cells being rare or not sufficiently deeply sequenced, the estimates may be unstable, and we recalibrate the estimates for this rare cluster to its most similar cluster to prevent potential false positives. Specifically, let *C*_*f* 1_index the rare group of cells; then, to identify the cell groups with the most similar open chromatin profile, we compute the correlation between 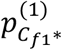 and 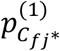 for all other clusters *j*= 1, …, *j*, across all regions. Assuming *C*_*f n*_ has the most similar chromatin profile, we rescale the current estimation of 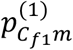 by the following formula:

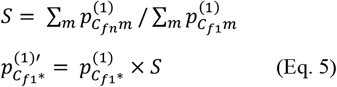

where S is the scale factor, 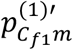 is the rescaled open probability estimate for the cluster *C*_*f* 1_ and feature *m*, and through rescaling, we are essentially assuming that most peaks are not differentially accessible between these two cell types.

### Cell type label prediction framework

Given a reference dataset, we estimate the probability of open chromatin 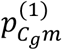 for each cell type *g* ∈ {1, …, *G*}, using the formula above. With a new set of observations 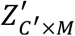, we apply the Bayes discriminative model to predict the corresponding cell type labels, 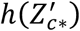.

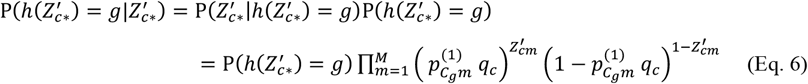

where 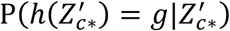 represents the posterior probability of cell 5 being sampled from cell group *g*, 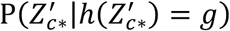 represents the conditional probability of observing 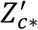 given that the cell *c* is sampled from cell type *g*, 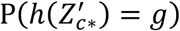 is the prior probability of a new observation belonging to cell group *g*, which can either be assumed to be a non-informative Dirichlet prior Dirich(δ) or estimated based on the cell type composition in reference data. Note that we have a large feature space so this choice will not make a big difference.

### Missing-corrected cumulative logistic regression (mcCLR)

Due to high sparsity of scATAC-seq data, perfect separability is common, hindering the parameter estimation in (Eq. 1). To address this issue, we incorporated Firth regularization (Eq. 2). Here we summarize the (unregularized) log likelihood function and information matrix for the cumulative response model and derive the analytical expression for the binary model. The loss function when considering cumulative response is

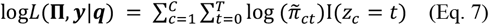

where C represent the total number of cells, π_*ct*_ and 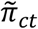 represent the probability of *t* PIC counts in cell *c* before and after accounting for cell-specific capturing probability, respectively. Specifically, π_*ct*_ = *P*(*Y*_*C*_ ≥*t*) − *P* (*Y*_*C*_ ≥ *t* + 1), Π_*C*_ = (π_*c0*_, π_*c1*_, π_*c*2_, …, π_*cT*_)^Trans^ and 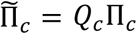, where *Q*_*c*_is the capturing probability matrix of dimension (*T*+ 1) × (*T* + 1) specified as

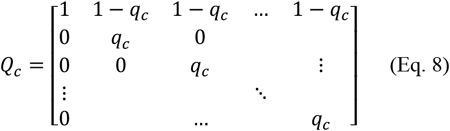

In our PACS model, an approximated estimation of parameters in the cumulative logit model were obtained using a method described in a previous set of studies^51,52^ that based on stacking the data and optimize with binary logistic regression specified by

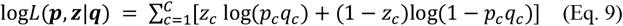

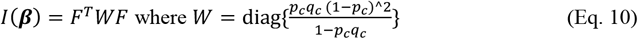

where *P*_*C*_ = *P* (*Z*_*C*_= 1).

### Parameter estimation for mcCLR

We implemented both Newton’s method and the Iterative Reweighted Least Squares method (IRLS) for parameter estimation. Briefly, for Newton’s method, ***β*** is estimated through the following iteration

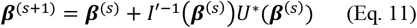

where the superscript *s* represents the iteration, *I ′* = *I* for the full model and *I* ′ = *I*_− {*d*}_ for the null model of β _{*d*}_ = 0. The score function *U*\*(*β*) is given by:

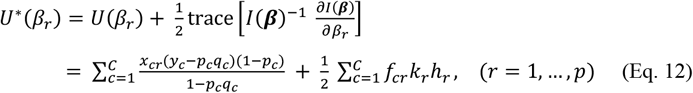

where the *h*_*C*_’s are the *c*^*th*^diagonal elements of the “hat” matrix, *H* = *W*^*1*/2^*F* (*F*^*T*^ *W F*) ^−1^ *F*^*T*^ *W*^*1*/2^, and 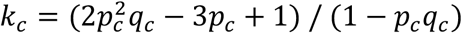.

For the IRLS method, the information matrix *I* is replaced with an estimate of the information matrix, 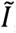,

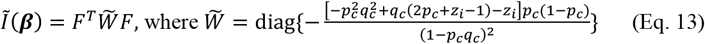

### Hypothesis testing framework of mcCLR

We utilized a generalized likelihood ratio test framework for hypothesis testing with the mcCLR model, although a Wald-type test can also be derived. As the model contains Firth regularization, we used the profile penalized likelihood approach to obtain *P* values^31,53^. Specifically, in the null model, the coefficients of interest are set to zero but still left in the model, so that the regularization accounts for the presence of these parameters during optimization.

### Data simulation for single factor differential test

To mimic real data, we estimated insertion rates 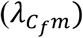 and *q*_*c*_ from the human cell line data and use these values to construct simulated data. Briefly, because viable snATAC-seq reads come from two adjacent Tn5 insertion events that have the right primer configuration (reviewed in^54^), we derived the size-filtered signed Poisson (ssPoisson) distribution to model this data generation process^24^. With the observed counts, we estimated the insertion rate parameters for two cell types, and regions with true insertion rate difference greater than 0.1 were set to be as true differential (H_a_) and the remaining region’s open probabilities were set equal (by taking the mean) and therefore non-differential (H_0_). Based on parametric model of latent and observed accessibility, we first sampled the latent ATAC reads by *ss*Poision 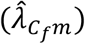 for *f* = 1,2, and then sampled the observing status by Bernoulli distribution parameterized by *q*_*c*_. The observed data were generated by the element-wise product of these two matrices. We randomly sampled 10,000 non-differential features to assess the type I error and 10,000 differential features to evaluate power. This simulation was conducted under varying numbers of cells in each group (from 250 to 1000), and each scenario was repeated 5 times.

### Data simulation for multi-factor differential test

Building upon the single factor setting, we further assumed the data to contain two cell types (T1 and T2) being sampled from two spatial locations (S1 and S2). We evaluated marginal effects and their interactions through separate simulations. To simulate data with marginal effect, the cell type effect is first introduced using the same method as in single-factor simulation, so the effect sizes varies. The spatial effect was then considered to affect features both with and without a cell type effect. Specifically, a third of the features with (and without) a cell type effect showed an accessibility difference across batches, with a log fold change of ±0.5. We introduced sample imbalance as frequently seen in real datasets. To be precise, we considered that S1 contained 1600 T1 cells and 800 T2 cells, while S2 contained 400 T1 cells and 1200 T2 cells. The peak by cell count data generation procedure is the same as for the single factor setting. To evaluate the performance of methods that do not support multi-factor testing, two strategies were used, naïve test and stratified test as reported in the main text. Following stratified test, we use Edington’s p value combination approach as it assumes consistent effect size across strata.

To evaluate interaction effect, we considered two configurations of interaction. In the first configuration, cells of T1 in S1 have high accessibility while other groups have low accessibility, with effect sizes also estimated from the cell line data. In the second configuration, cells of T1 in S1 and cells of T2 in S2 have high accessibility while other groups have low accessibility. The second configuration may not be common in real biological data, but a method with capability of testing interaction effect should be able to identify it.

### Data simulation for time-series differential test

To evaluate model performance in situations where the design matrix contains ordinal covariates, we simulated time-series snATAC-seq data across five time points. We assumed linear and quadratic temporal effects on accessibility and set the effect size (log fold change) to be 0.3 or 0.5 between the two groups. The baseline accessibility was generated from the cell line data and the peak by cell count data generation procedure is the same as for the single factor setting. N=1000 cells were sampled for each time point.

### Evaluating type I error and power in real datasets

To estimate type I error in real data where the ground truth is unknown, we used a label permutation approach, where the data in one cell type were divided randomly into two groups, and a differential test was conducted between these groups. As this is randomly assigned, all features were believed to be non-DAR, so the proportion of *P* values smaller than 0.05 is the empirical type I error using real data. Then, we set the fifth rank percentile as the correct critical value for those methods with type I errors greater than 0.05. We next conducted a test with two different cell types using the calibrated critical values for each method. Since we do not know the true DAR set, we defined the pseudo-true DAR peaks as the union DAR set of all tested methods, using their corresponding new critical values. Power for each method was then calculated by the number of DARs detected divided by the number of pseudo-true DARs. This approach is adopted from Ref.^24^.

### Estimating effect size (fold change and accessibility change)

A common practice to determine differential features in single cell data is by setting a cutoff for both *P* value and fold change. In scRNA-seq data analysis, one way to estimate the effect size of a particular variable (predictor) is by calculating the fold change (FC) for the normalized data, obtained by dividing the normalized mean expression of one group by the other group. However, with snATAC-seq data, there is no direct normalization method available, and computing the fold change on raw read counts may lead to inaccuracies due to disparities in data capture. Here, we propose to use the capturing probability-adjusted count to compute fold change (FC) or the arithmetic difference between accessibility (accessibility change, AC) of two cell types. To be precise:

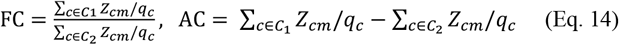

where *m* is the feature of interest and *C*_1_ and *C*_2_ are the lists of cells that contain foreground and background cell types.

### Processing kidney adult data with Signac

We used Signac^23^ to evaluate the effectiveness of our method in correcting for batch effect at the feature level. We follow the standard workflow as recommended in the Signac vignette (https://stuartlab.org/signac/articles/pbmc_vignette.html). Briefly, we used the TF-IDF approach without feature selection (*min*.*cutoff = ‘q0’*), followed by SVD to reduce dimensionality. We then conduct clustering and UMAP visualization using the dimensions 2 to 30 (as the first LSI dimension usually reflects sequencing depth, per the Seurat tutorial). The sample and cell type labels are retrieved from the annotations in the initial publication.

### Batch mixing score calculation

We calculated the batch mixing scores in the PCA space as a measure of batch effect. At the cell level, the batch mixing score is adopted from Ref.^38^ and is defined as the proportion of nearest neighbor cells with different batch identities, where a higher score indicates better mixing between batches, and thus a smaller batch effect. At the whole data level, the batch mixing score is defined as the mean batch mixing score across all cells. To calculate the expected batch mixing score for a given dataset when no batch effect is present, let *M* denote a cell type-by-batch matrix, with each element *m*_*ij*_ representing the number of cells in the cell type *i* and batch *j*. Then the expected data-level batch mixing score in the setting of no batch effect is given by

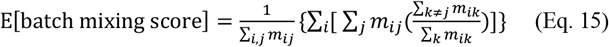

The normalized batch mixing score is the batch mixing score divided by the expected score under random mixing, and thus a higher normalized batch mixing score indicates better mixing across samples.

### Processing developing human brain data

This dataset contains 18 specimens collected from human donors. For our study, we excluded samples with unknown spatial locations (GW17, GW18, GW21) or samples not from the cortex (MGE_GW20 and MGE_twin34). Here we focused on the excitatory neuron lineage, including radial glia (RG), intermediate progenitor cells (IPCs), early excitatory neurons (earlyEN), deep layer excitatory neurons (dlENs), and upper layer excitatory neurons (ulENs). We further excluded the insular region for having too few cell counts (645 cells across five cell types). The data matrix was saved as a binary matrix, so we implemented the missing-corrected logistic regression model for the analyses of this data.

### DAR identification in the developing human brain data

We constructed two models to identify the significant region effect of the excitatory neuron lineage. Specifically, to identify the region effect, the systematic component of the PACS model is specified as:

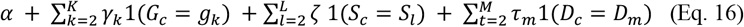

where *G* is the index of cell type, *S* is the index of spatial location, and *D* is the index of the donor. The null hypothesis for the test is *H*_0_: ζ = 0. To identify the cell type specific region effect, we additionally included the interaction terms between each cell type and spatial location, and the test was conducted for each interaction term.

### Motif enrichment analysis

The motif enrichment analysis was conducted with Homer^55^. The list of significant DAR peaks is used as input for the analysis, with the size of the search region specified as 300 bp around the peak center. The reported motif enrichment scores are FDR-corrected *P* values from the known motif results.

### DAR identification in the human PBMC treatment data

To identify the cell type-specific temporal effect in the PBMC treatment data, the systematic component of the PCAS model is specified as:

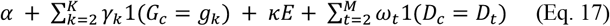

where *G* is the index of cell type, *E* is the experimental time index (0, 1, 2 corresponds to control, 1h, and 6h after treatment, respectively), and *D* is the donor index. The null hypothesis for the test is *H*_*0*_: *κ* = 0.

### Gene and pathway enrichment with GREAT

We used the GREAT method (v. 4.0.4) to conduct gene and enrichment analysis^39^, with DARs as input and default parameter settings. The output from GREAT for the human PBMC data can be found in the **Supplementary Tables 17-18**.

## Supplementary Materials

Supplementary Figures 1-5

Supplementary Table 1: Parameter estimation using simulated data

Supplementary Table 2: Type 1 error and power of different methods using simulated data (one-factor setting)

Supplementary Table 3. Type 1 error and power of different methods using simulated data (two-factor setting)

Supplementary Table 4. PCT specific peaks in the adult kidney data

Supplementary Table 5. PST specific peaks in the adult kidney data

Supplementary Table 6. GREAT gene enrichment results of PCT specific peaks

Supplementary Table 7. GREAT gene enrichment results of PST specific peaks

Supplementary Table 8. Number of cells in across spatial regions and donors

Supplementary Table 9. V1 specific peaks in the developing human brain data

Supplementary Table 10. PFC specific peaks in the developing human brain data

Supplementary Table 11. Homer motif enrichment results of the V1 region in the human developing brain data

Supplementary Table 12. Homer motif enrichment results of the PFC region in the human developing brain data

Supplementary Table 13. Number of differential peaks between PFC and V1 across excitatory neuron lineage in the developing human brain data

Supplementary Table 14. Significant up-regulated peaks after treatment across cell types in the PBMC treatment data

Supplementary Table 15. Significant down-regulated peaks after treatment across cell types in the PBMC treatment data

Supplementary Table 16. Number of significant differential peaks after treatment across five cell types, using PACS or ArchR

Supplementary Table 17. GREAT pathway enrichment results of up-regulated treatment effect peaks in T cells

Supplementary Table 18. GREAT gene enrichment results of up-regulated treatment effect peaks in T cells

## Code Availability

PACS is an open-access software available at the GitHub repository https://github.com/Zhen-Miao/PACS. Codes for reproducing the analyses are also available at the GitHub page.

## Author Contribution

JK and ZM conceived the study. ZM, JW, and JK designed the statistical model. JW formulated the missing data model for sequencing depth and derived the analytical expression for missing-corrected logistic regression estimation procedure. ZM implemented the model and constructed the software package with feedback from JW, DK, and JK. ZM conducted the simulation and real data analysis with help from KP and DK. JK supervised the work. JK and ZM wrote the manuscript with feedback from JW.

## Acknowledgements

This work has been supported in part by the UC2DK126024 and R01 DK126925-05 grant to JK and also by the Health Research Formula Fund of the Commonwealth of Pennsylvania who did not play a direct role in the work. We thank Blavatnik Family Fellowship that supported the work of ZM. We thank Dr. Pablo Camara, Dr. Nancy Zhang, Dr. Kui Wang, Dr. Xiangjie Li, Dr. Yinan Lin, Dr. Mengying You and members of Junhyong Kim’s lab, especially Dr. Erik Nordgren for their constructive suggestions that improved this work. We thank Dr. Kun Zhang and Dr. Jason Buenrostro for sharing the metadata.

## Competing interests

The authors declare no competing interest.

## Notes

### Competing Interest Statement

The authors have declared no competing interest.

### Summary of Updates

Manuscript title corrected; Figure 3 revised; Supplementary figures re-organized; Grant information updated

