## Supplementary Figures for "PACS allows comprehensive dissection of multiple factors governing chromatin accessibility from snATAC-seq data"

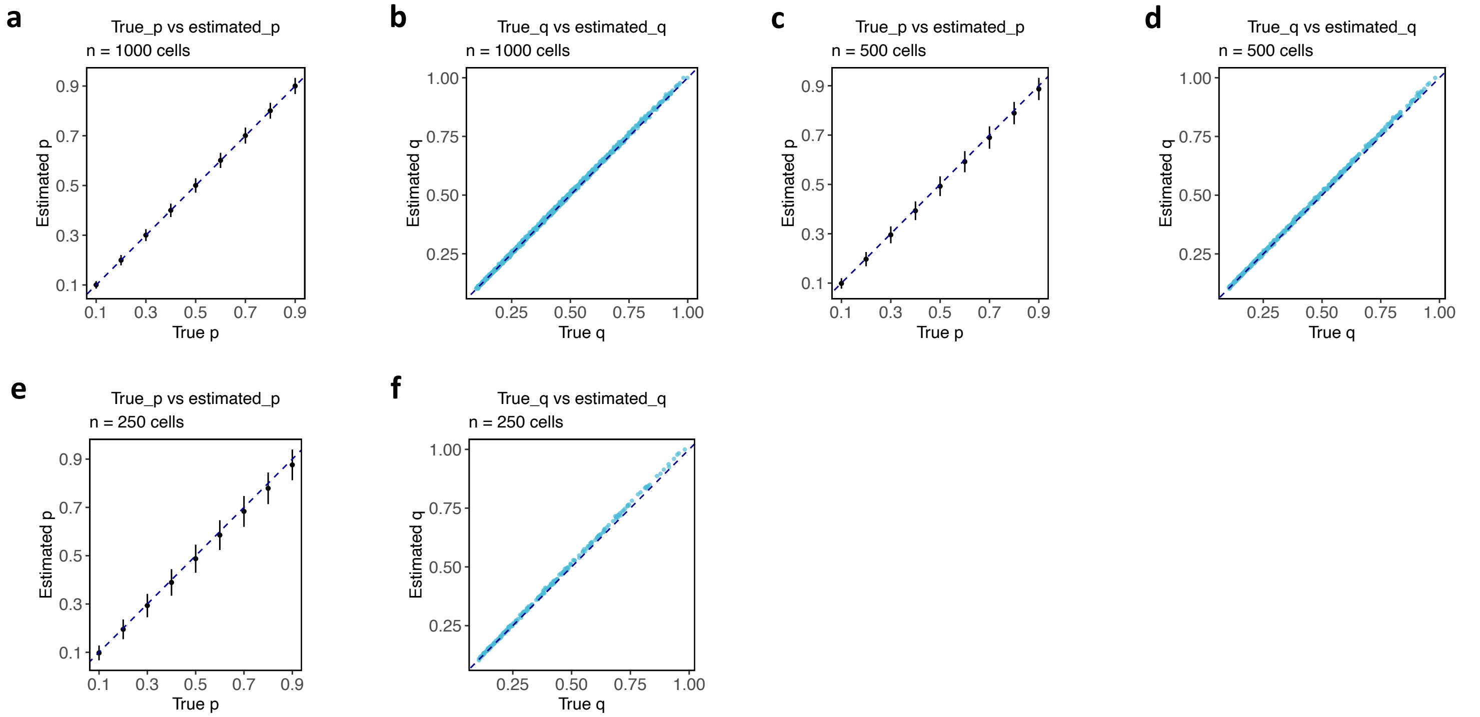


**Figure 1. Parameter estimation evaluation and application to cell type annotations.**

**a-f**. Parameter estimation accuracy evaluated using simulation data, for n=1000 cells (**a-b**), n=500 cells (**c-d**) or n=250 cells (**e-f**). Here p represents P(y >= 1) and q represents the capturing probability. The error bars indicate the standard deviation across repeated simulations (n=5).

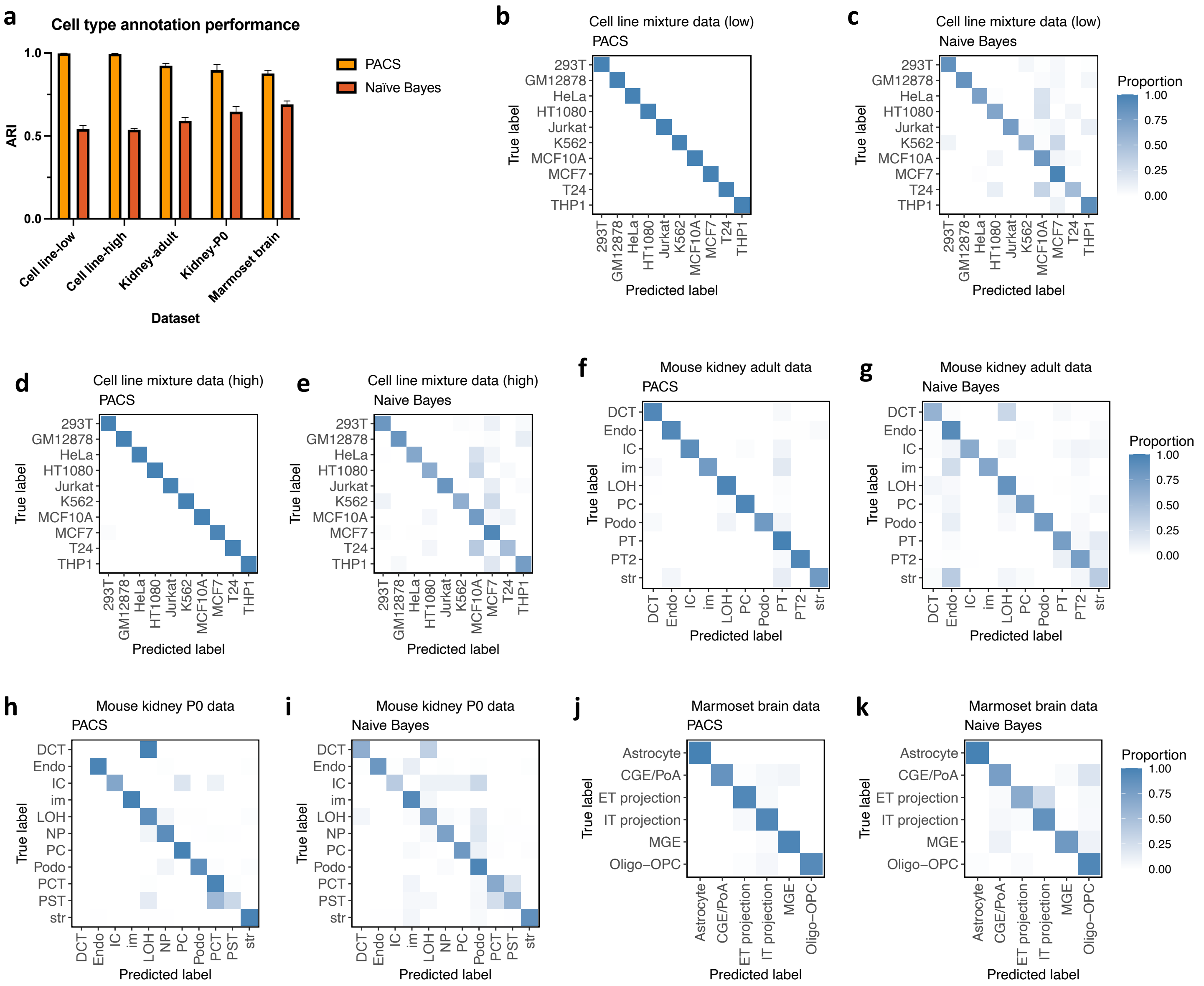


**Supplementary Figure 2.**

**a**. Comparison of cell type annotation adjusted rand index (ARI) between PACS and Naïve Bayes method for five datasets.

**b-k**. Confusion matrix between true cell type labels and PACS-inferred (or Naïve Bayes-inferred) cell type labels for five real datasets.

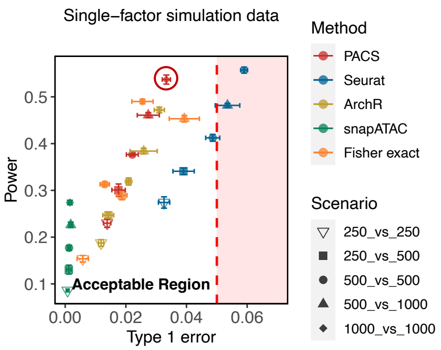


**Supplementary Figure 3.**

Type I error and power of different methods evaluated with single-factor simulation data. The tests are two-sided, and the error bars indicate the standard deviation across repeated simulations (n=5). The best-performing method is indicated by a circle with matched color.

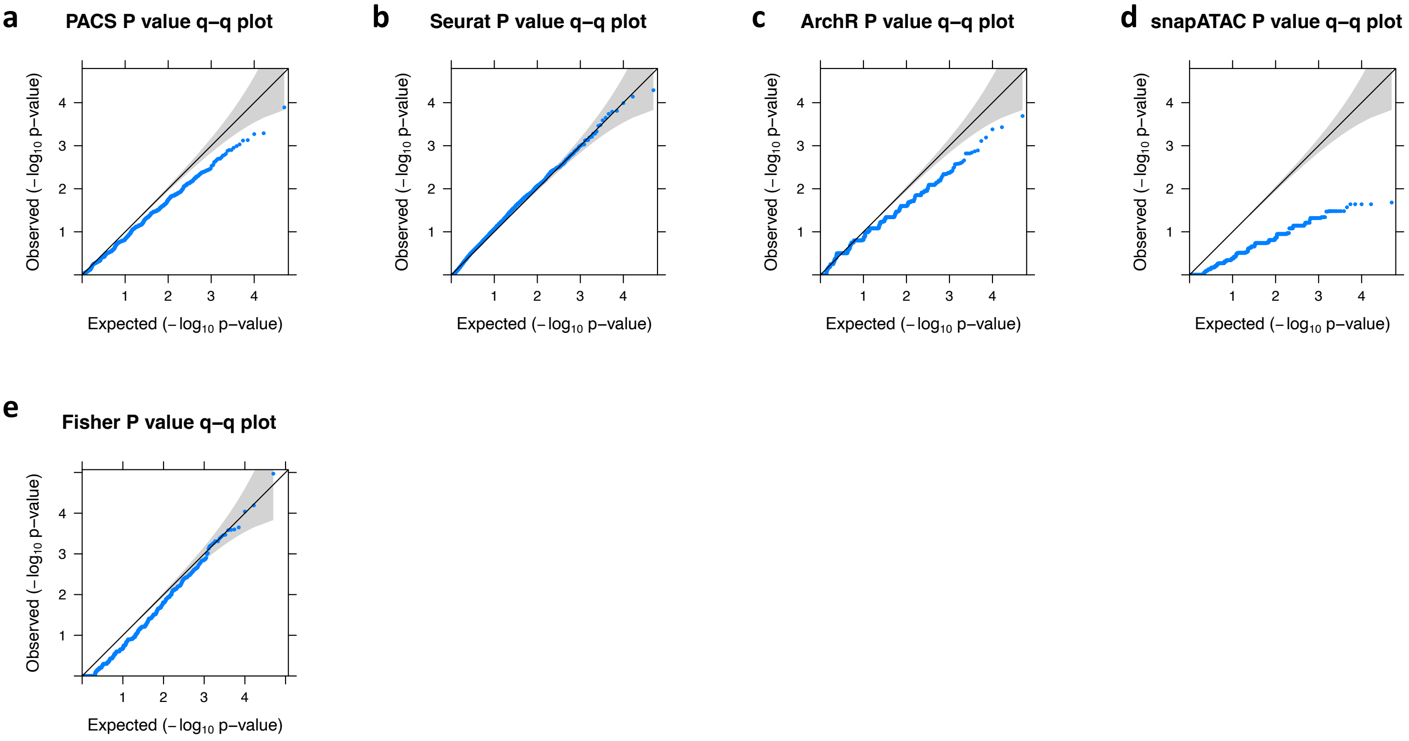


**Supplementary Figure 4.**

**a-e**. Quantile-quantile plots for P values under the null for five testing methods, using simulated data with no insertion rate difference.

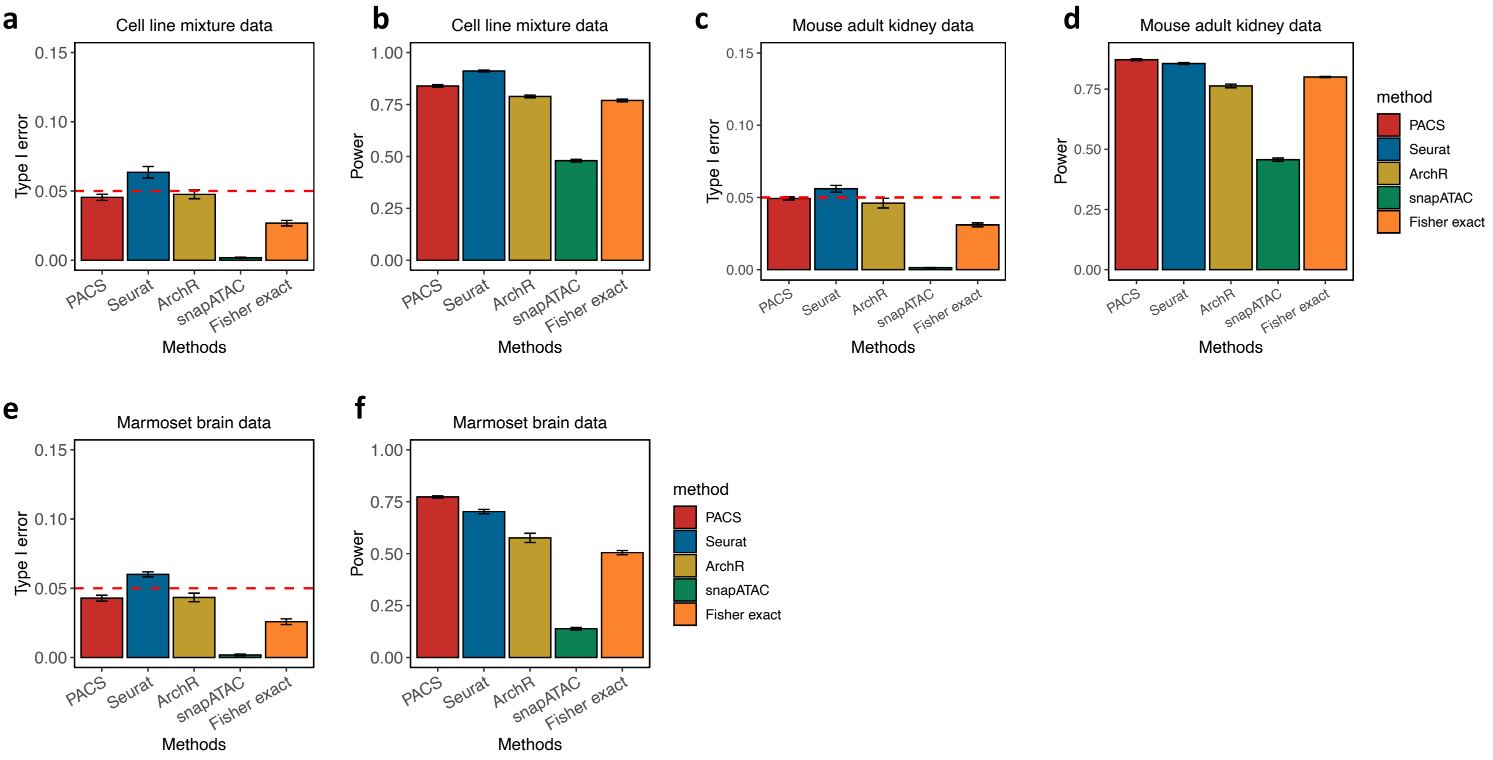


**Supplementary Figure 5.**

**a-f**. Type I error and power of different statistical testing methods evaluated with empirical cell line mixture data (**a-b**), adult mouse kidney data (**c-d**), or marmoset brain data (**e-f**). N=1000 cells were sampled for each group (cell type).

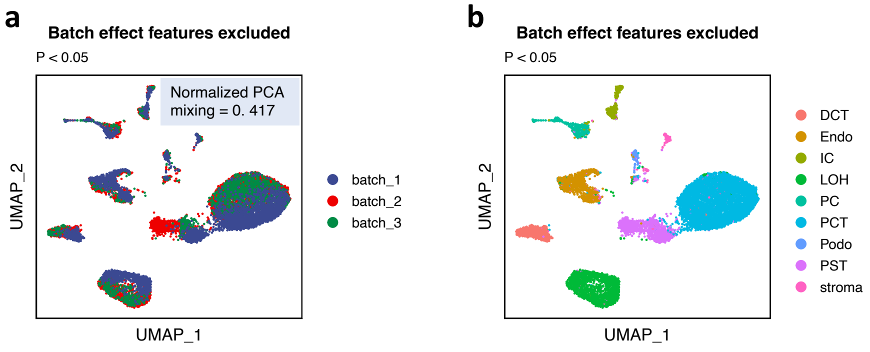


**Supplementary Figure 6.**

**a-b**. UMAP dimension reduction plot constructed after excluding features with significant batch effect (P value < 0.05, no FDR correction), colored by batch labels (**a**) or cell types (**b**). Features with batch effect are detected with PACS differential test module. Normalized PCA mixing represents the normalized mixing score calculated in the PCA space, with 1 being no batch effect and 0 being strongest batch effect.

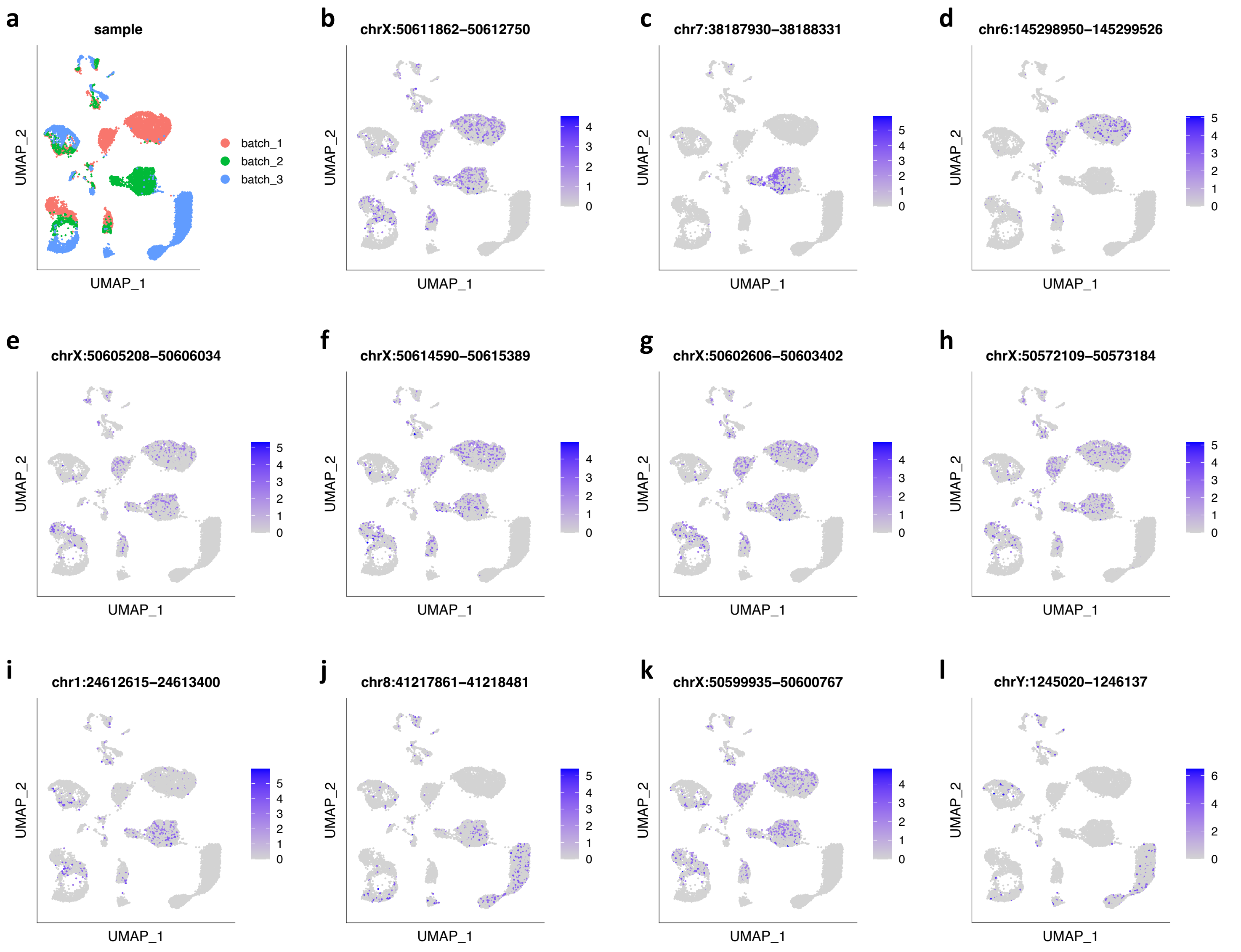


**Supplementary Figure 7.**

**a**. UMAP dimension reduction plot constructed with all features, colored by batch labels. This panel is identical to **Fig. 4a**, and is displayed here for examining feature plots in panels **b-l**.

**b-l**. Feature plots for top significant batch effect peaks determined by PACS.

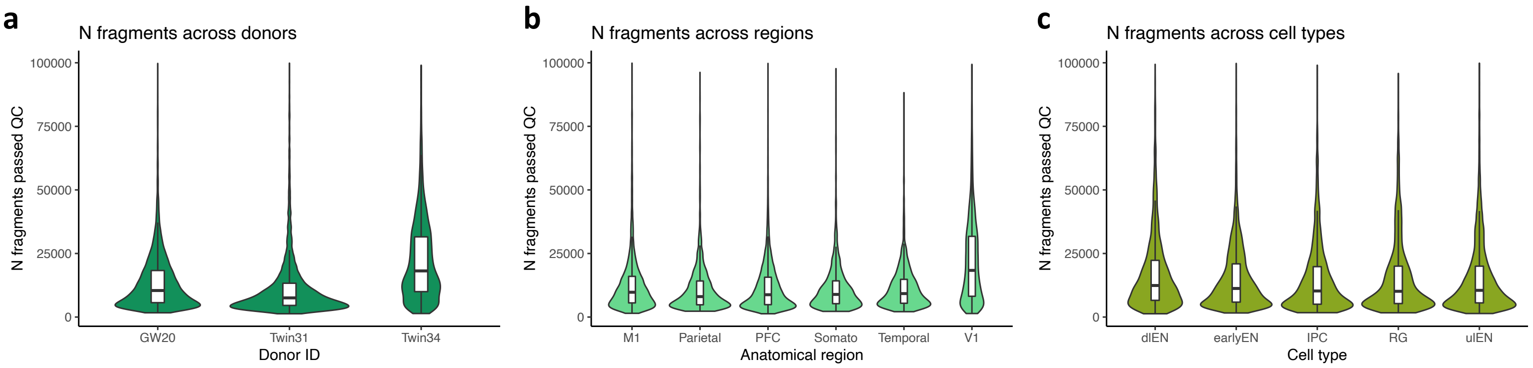


**Supplementary Figure 8.**

**a-c**. Violin plots that summarize number of fragments in each cell across different donors (a), brain regions (b), or cell types (c), for the human brain data. Center line in box plot represents median and the lower and upper hinges correspond to the first and third quartiles. The upper or lower whisker corresponds to 1.5 times the inter-quartile range or the largest/smallest values.
